# Density-Dependence and agricultural intensity affect host-parasite-parasitoid interactions in a bird-insect trophic system

**DOI:** 10.64898/2026.10.06.757064

**Authors:** Simon Coroller-Chouraki, Jade Savage, Marc Bélisle

## Abstract

Agricultural intensification is a major cause of population declines in farmland birds and insects worldwide. While these declines are expected to disrupt trophic interactions through density-dependent cascading effects, examples outside of crop pest control or involving more than two trophic levels remain poorly documented. Here we quantify such cascading effects across three trophic levels involving a declining aerial insectivore, the Tree Swallow (*Tachycineta bicolor*), which breeding success decreases with agricultural intensity, *Protocalliphora* bird blowflies (Diptera: Calliphoridae) that parasitize its nestlings, and *Nasonia* parasitoid jewel wasps (Hymenoptera: Pteromalidae) that feed upon blowfly pupae within swallow nests. Agriculture intensity may affect each of these actors either directly via, for instance, toxicological effects from pesticide use, or indirectly through density-dependent trophic interactions, which can exacerbate or mitigate one another. Using Bayesian models within a causal inference framework, we quantified how nestling host density, meteorological conditions, and agricultural landscape context affect *Nasonia* parasitoid infestation of *Protocalliphora* pupae, and estimated *Protocalliphora* emergence success. Analyses were based on 3,340 *Protocalliphora* puparia collected from 248 Tree Swallow nests over a 16-year period (2004-2019) across a 10,200-km^2^ agricultural intensity gradient in Southern Québec, Canada. Model predictions indicated that *Protocalliphora* emergence probability increased steeply with nestling host availability (*i.e*., nestling-days). Whereas *Nasonia* parasitoid infestation occurrence at the nest level increased with *Protocalliphora* host availability, the proportion of infested *Protocalliphora* pupae within nest decreased and led to a dilution effect at the individual level. Although *Protocalliphora* abundance within nests decreased with agricultural intensity, their emergence probability peaked in agrointensive landscapes. This increase was associated with reduced *Nasonia* infestation rates, suggesting that agricultural intensity indirectly mitigated parasitoid pressure on *Protocalliphora* host populations. Our results highlight how agricultural intensification can shape multitrophic interactions and life- history traits (emergence success and infestation rates) by simultaneously reducing host availability and weakening top-down control by parasitoids.

## Introduction

The composition, structure, and dynamics of trophic networks, including their stability and resilience, can be strongly shaped by density-dependent processes (Polis and Strong 1996, Polazzo et al. 2022, 2023, Ulanowicz et al. 2014). For instance, variation in host density, social behavior, and spatial clustering can noticeably influence parasite (*sensu lato*) diversity, transmission, and prevalence (Patterson and Ruckstuhl 2013, Wells et al. 2015, Gunton and Pöyry 2016). Predicting when and how such processes will affect trophic networks is, however, challenging given that the form and strength of density-dependence has been found to be highly nonlinear and context-dependent. For example, density-dependence and its consequences (e.g., carry-over and stabilizing effects) for populations composing trophic networks can vary across trophic levels and life-stages (Hellriegel 2000, Ratikainen et al. 2008). They have also been found to vary across spatial scales and environmental conditions, including landscape and habitat heterogeneity and patchiness (Hanski 1999, Hassell 2000, Abrams and Vos 2003, Lebreton 2009, Gunton and Pöyry 2016). The diversity and context-dependence of density-dependent mechanisms and responses that may occur flares out in multi-trophic systems in which direct and indirect effects among interacting species can exacerbate, dampen, or even cancel themselves. Because the composition, structure, and dynamics of trophic networks have been studied predominantly in simplified models or laboratory systems, additional field studies encompassing multiple interacting species are needed to assess and improve the generality of current understanding.

For instance, the consequences of agricultural intensification for trophic interactions have been extensively studied in biological control systems involving crops, insect pests, and their natural enemies (Agrawal 2000, Gagic et al. 2012, Tooker and Pearsons 2021). However, this body of work has focused primarily on relatively simple food webs involving a limited number of target species within highly managed environments, such as greenhouses, or has typically been restricted to small spatial scales. Consequently, whether the ecological principles emerging from these systems apply to natural multitrophic communities distributed across heterogeneous agricultural landscapes remains largely unresolved. Addressing this question is critical because agricultural intensification can simultaneously alter landscape structure (i.e., habitat composition and configuration), and thus landscape heterogeneity, microclimatic conditions, the phenology and availability of both resources and consumers, as well as pesticide exposure (e.g., Stanton et al. 2018, Wagner 2020, Wagner et al. 2021). These changes can influence multiple trophic levels through both direct environmental effects and indirect trophic pathways (Halsch et al 2026). By modifying the topology and strength of trophic interactions, and potentially disrupting trophic regulation, these perturbations can generate cascading and knock-on effects that propagate throughout ecological networks (Abrams and Vos 2003, Ripple et al. 2016, Eisenhauer et al. 2019, Polazzo et al. 2023). Pesticides are particularly problematic in this context as they often affect non-target organisms, including natural enemies, more severely than their target pests, thereby creating trophic imbalances that promote pest outbreaks (Geiger et al. 2010, Barbosa et al. 2012, Janssen and van Rijn 2021). Quantifying cascading effects has thus become increasingly urgent given the potential for enemy release that can exacerbate impacts on biodiversity or impair ecosystem services and the widespread and concurrent population declines of insects and insectivorous birds across agricultural landscapes (Kehoe et al. 2021, Wagner et al. 2021, Douglas et al. 2023, Rigal et al. 2023).

Bird blowflies of the genus *Protocalliphora* (Diptera: Calliphoridae) provide an excellent model for investigating cascading trophic effects in agroecosystems. Species of this genus are widespread in bird nests throughout the Holarctic, where their larvae are obligate hematophagous ectoparasites of nestlings, including those of farmland birds (Lehmann 1993, Nogueira et al. 2023). Following pupation, *Protocalliphora* pupae become hosts for parasitoid wasps of the genus *Nasonia* (Hymenoptera: Pteromalidae), forming a naturally occurring tri-trophic network (Daoust et al. 2012a). In cavity-nesting birds such as the Tree Swallow (*Tachycineta bicolor*), the number and size of *Protocalliphora* puparia recovered from nests provides a reliable proxy for local parasite abundance and performance (Jánošková et al. 2010, Coroller-Chouraki et al. 2026b).

Recent work has shown that agricultural intensification indirectly reduces *Protocalliphora* prevalence and infestation intensity in Tree Swallow nests by decreasing swallow abundance at both the landscape (breeding density) and nest (clutch or brood size) scales, thereby limiting host availability for these bird blowflies (Coroller-Chouraki et al. 2026b). At the highest trophic level, *Nasonia* parasitoids (which can kill up to 90% of *Protocalliphora* pupae in a given year; Coroller-Chouraki et al. 2026a) are also likely to experience the impact of a reduced availability of suitable hosts (Hassell 2000, Skovgård and Nachman 2004, Aung et al. 2011). Moreover, these parasitoids are especially susceptible to contamination by pesticides, which can impair their survival, host-searching behavior, and reproductive success (Sanchez-Bayo and Goka 2014, Pisa et al. 2015, Schöfer et al. 2023). Agricultural intensification may therefore reshape this model tri-trophic system through a combination of direct effects on individual trophic levels and indirect bottom-up and top-down pathways that alter the strength of trophic interactions.

Disentangling the direct and indirect effects that may be at play in a multi-trophic network can however be challenging as several to many confounding relationships may bias estimates of the strength of trophic interactions. To reduce such biases, we apply a causal inference framework to 16 years of nest-box monitoring data collected across a 10,200 km² agricultural intensity gradient in southeastern Canada to investigate how agriculture may differentially affect the probability that *Protocalliphora* ectoparasites of Tree Swallow nestlings are attacked by *Nasonia* parasitoids, and in the event that they escaped parasitoidism, their probability of emergence. Building on previous evidence that agrointensive landscapes support lower nestling host availability and reduced *Protocalliphora* prevalence and loads in swallow broods (Coroller-Chouraki et al. 2026b), we tested three hypotheses. First, because parasitoids are particularly sensitive to agricultural intensification through both reduced host availability and anthropogenic impacts such as those resulting from pesticide use (Jonsson et al. 2012, Syahidah et al. 2021, Molina et al 2019), we predicted that *Nasonia* parasitoidism would decline with decreasing *Protocalliphora* loads in swallow broods and increasing agricultural intensity after controlling for the latter effect. Second, given the generally harsher conditions experienced by insects in agrointensive landscapes (Halsch et al. 2026, Seimandi-Corda et al. 2026), we predicted that the emergence success of *Protocalliphora* pupae that escaped *Nasonia* parasitoidism would also decrease with agricultural intensity. Third, because, when facing a perturbation, populations of species occupying higher trophic levels tend to decline more strongly than those from lower levels despite context-dependent responses (Gagic et al. 2012, Zhao et al. 2015, Wagner 2020), we predicted that reduced *Nasonia* parasitoidism in agrointensive landscapes could partially offset the negative effects caused by agriculture on *Protocalliphora* and thereby lead to improved *Protocalliphora* overall emergence success.

## Methods

### Study system

The study system was established in Quebec, Canada, in 2004 (Ghilain and Bélisle 2008) and covers approximately 10,200 km². It comprised 400 nest boxes distributed equally among 40 farms, located along an east-west gradient of agricultural intensity (Fig. 1). All nest boxes were placed ∼1.5 m above ground and spaced ∼50 m apart along a linear transect that followed field edges. Eastern agricultural landscapes are dominated by pastures and forage crops embedded into substantial forest while western ones consist primarily of large expanses of annual row crop monocultures mostly focused on maize, soybean, and wheat associated with heavy pesticide use (Giroux 2022).

**Figure 1.**
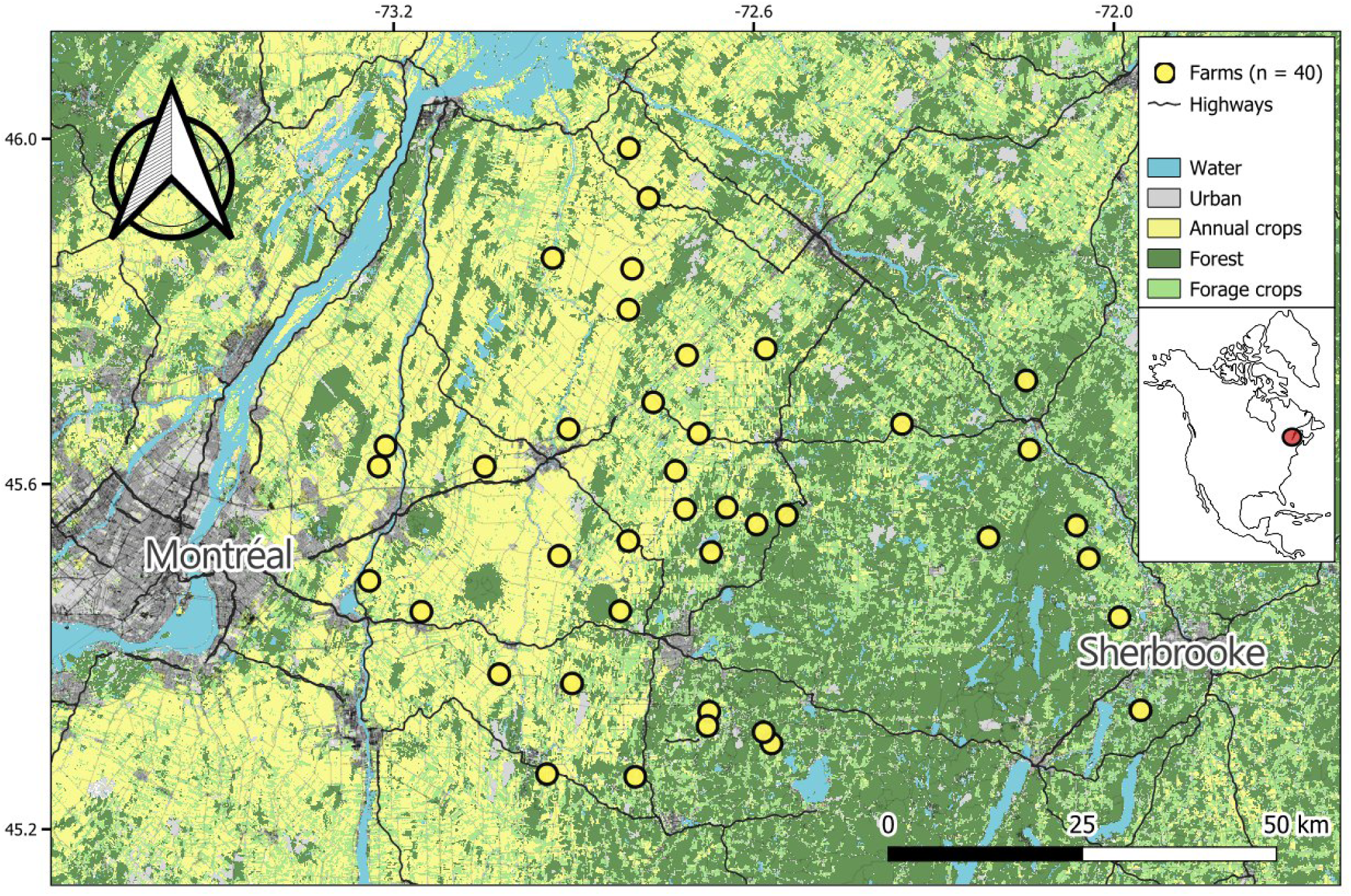
The distribution of the 40 farms across a 10,000 km² area in South-Eastern Canada (QC), representing a gradient of agricultural intensification. Coordinates are in the Universal Transverse Mercator (UTM) zone 18 north, while the mapping was conducted using the NAD83 MTM zone 8 system.

To build causally unbiased models and accurately predict how agricultural landscapes, meteorology, avian host density, and *Protocalliphora* ectoparasite density affect both emergence and infestation rates, we gathered nest-specific environmental data and controlled for these factors statistically (Cinelli et al. 2022).

### Agricultural landscape characterization

We surveyed landscape composition within a 500m radius of the 400 nesting boxes at the end of each swallow breeding season between 2006 and 2019. Habitat types were categorized “Forest”, “Corn/Soybean”, “Other Cereals” (i.e., wheat, oat, barley), and “Pasture/Forage” (i.e., hay, alfalfa, clover). We used orthophotos (scale 1:40,000) and QGIS software to delineate habitat boundaries estimating habitat coverage within the 500m radius. Following methods by (Garrett *et al*. 2022a, Powell *et al*. 2024), we performed a robust compositional Principal Component Analysis (rPCA) using the *robCompositions* R package (v2.4.6; Templ *et al*. 2011) on the yearly relative areas covered by the above habitat types around each nest box. Streams and water bodies covered <1% of nest box surroundings (Garrett *et al*. 2022b) and were not considered. The first rPCA axis (Comp.1) explained 78.99% of landscape variance, being positively correlated with maize and soybean cover and negatively with forage and trees. The second axis (Comp.2) explained 16.65% of the variance, correlating positively with trees and negatively with forage.

Because crop cover data were missing in 2004–2005, Comp.1 and Comp.2 scores were extrapolated by averaging 2006–2008 values for each nest box, as landscape composition varied little among years (see Fig. S1). This ensured each nest box was assigned yearly landscape scores, which have been shown to influence Tree Swallow phenology (Bourret *et al*. 2015, Courtois *et al*. 2021) and breeding success (Garrett *et al*. 2022a,b,c). We used the first two components values fort each nest box as landscape predictor variables to model and statistically control for the effects of the agricultural contexts.

### Meteorological variables

We extracted hourly temperature (°C) and precipitation (mm) values based on the GPS coordinates of each farm from 2004 to 2019 using the *NASAPOWER* package in RStudio (v4.3.1; Sparks 2024), which provides hourly climatic data with a resolution of 50km*50km from NASA’s Prediction of Worldwide Energy Resource (POWER) project. For each individual nest, we filtered temperature and precipitation to cover the period from the first hatching date to seven days after the last nestling fledged or died. This period was chosen to encompass the full development and pupation period of *Protocalliphora* spp. (Coroller-Chouraki *et al*. 2026b). The hourly data were then averaged across these periods to obtain the mean temperature and mean precipitation in a given nest. We used these values as explanatory variables to control for mean meteorological contexts during which *Protocalliphora* develop in each nest.

### Biological data acquisition

Tree Swallows nests were monitored every two days during the breeding season to assess clutch size, nestling survival, or fledging age between 2004 and 2019. We defined ‘phenology’ as the timing of potential *Protocalliphora* access to nestling blood, approximated by the date of the first hatching event for each clutch. To assess ‘nestling availability’ for *Protocalliphora* ectoparasites, we summed up the number of nestling-days for each nest. A previous study on this system showed that ‘nestling availability’ was a strong predictor of *Protocalliphora* occurrence and abundance (Coroller-Chouraki *et al*. 2026b).

At the end of each breeding season, we collected and bagged the entire content of nest boxes exclusively occupied by Tree Swallows where we had detected at least one hatching event. Collected nests were stored at 4°C after 2 days at −80°C to kill any remaining living organisms, before processing. All nest material was sorted under a ventilated hood. All *Protocalliphora* spp. puparia found within were counted and stored in vials filled with 75% ethanol and each was referenced according to farm and nestbox IDs as well as year of sampling. Note that puparia from 2008 and 2009 were collected under another protocol by Daoust *et al*. (2012b) to rear some material in the lab and collect emerging flies or *Nasonia* spp. wasps; those years were therefore excluded from our analysis.

Given the considerable number of *Protocalliphora* and the time required for examination, we analyzed a subsample of 248 nests. First, ten farms were selected using stratified random sampling to represent the regional gradient of agricultural intensity between 2004 and 2019. For each year (except 2008 and 2009), two infested nests (when available) were randomly sampled from each of these ten farms. All *Protocalliphora* puparia found within these nests were analyzed. When fewer than 100 puparia were available for a given year, additional farms were randomly added (each contributing up to two infested nests) until an annual minimum of 100 puparia was reached. On average, 1.2 additional farms per year were included in the subsamples, a number unlikely to alter general patterns derived from the core group of ten farms.

In total, 3,340 *Protocalliphora* puparia from 248 nests were examined. Each puparium was observed under a stereomicroscope, and when preservation allowed, identified to species using Sabrosky *et al*. 1989, as well as Whitworth 2019. All puparia were inspected for signs of*Nasonia* parasitoidism, determined from characteristic 0.5-1 mm exit holes or by the presence of unemerged larvae within the puparia (Coroller-Chouraki *et al*. 2026a). For well-preserved and unparasitized puparia, emergence success was assessed from the presence of the emergence cap and residual pupal remains.

### Directed Acyclic Graph and Statistical Analysis

Hierarchical models built within a causal inference framework were used to assess the influence of various environmental parameters on the probability of *Nasonia* infestation in *Protocalliphora* puparia, and *Protocalliphora* subsequent probability of emergence. As *Protocalliphora* pupae infested by *Nasonia* wasps die and therefore do not emerge, we modeled all uninfested *Protocalliphora* pupae to investigate the agroenvironmental factors affecting emergence probability, excluding parasitoidism. We used a causal inference framework to isolate direct and indirect effects while controlling for confounding variables (Cinelli *et al*. 2022). For instance, a higher density of *Protocalliphora* in nests could affect the likelihood of *Nasonia* parasitoids infesting them, leading to an emergence failure, while also having other density-dependent influence on the emergence success of uninfested pupae (Fig. 2).

**Figure 2.**
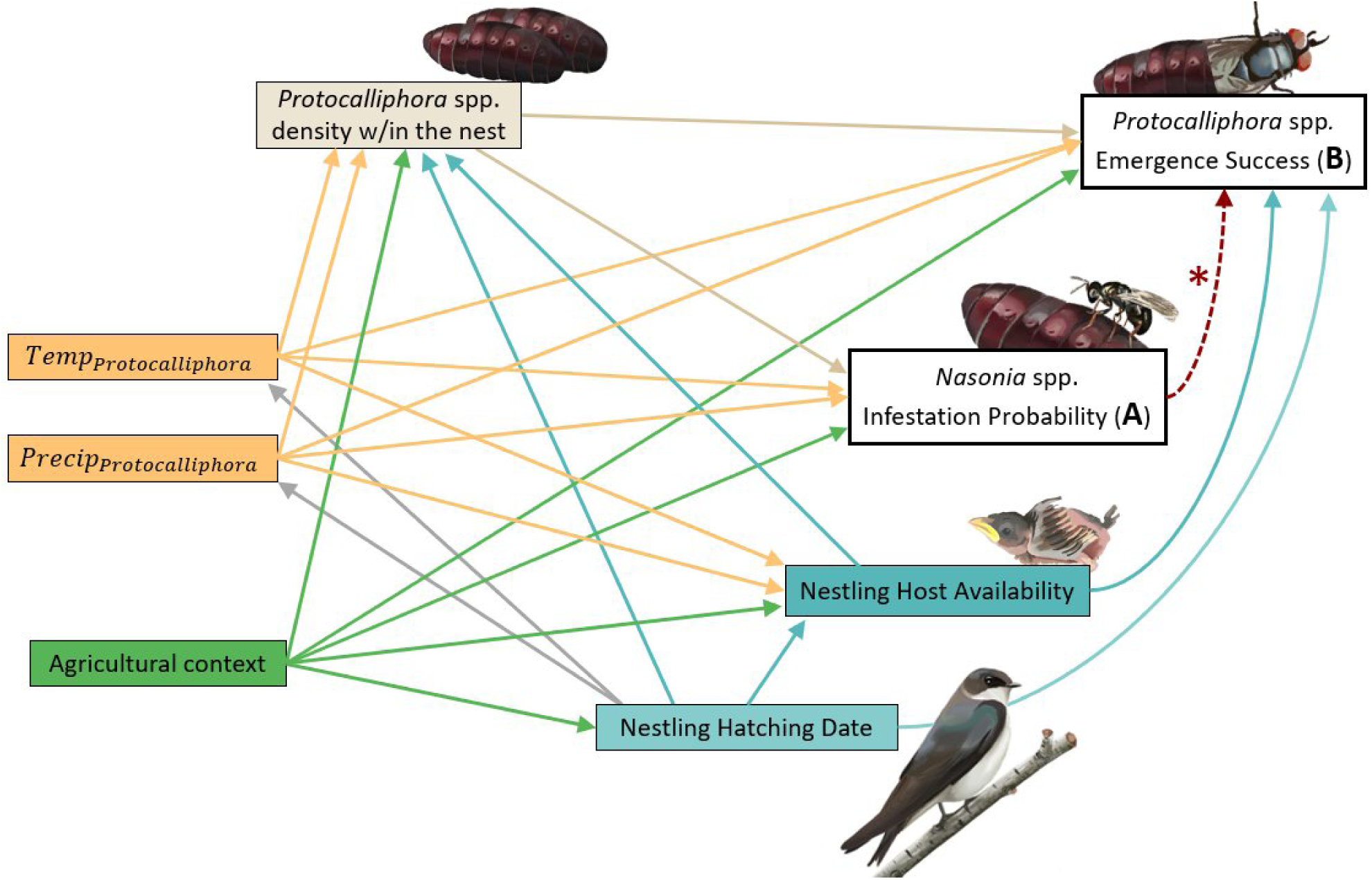
Directed Acyclic Graph of putative causal relationships among studied variables. Direct effects of variables on (A) the effects of environmental factors on the probability of *Nasonia* infestation in *Protocalliphora* puparia, and (B) the effects of environmental factors on the emergence probability in uninfested *Protocalliphora* puparia (see models in Table 1). * The red dotted arrow indicates a deterministic biological constraint: infestation by *Nasonia* spp. systematically leads to *Protocalliphora* mortality, resulting in zero emergence success (perfect separation). To identify environmental drivers, emergence success was first modeled using only uninfested puparia (B1; Table 1). A second model was then performed at the population scale, including all puparia (infested and uninfested), to assess the overall impact of parasitoidism on emergence probability (B2; Table 1).

**Table 1.**
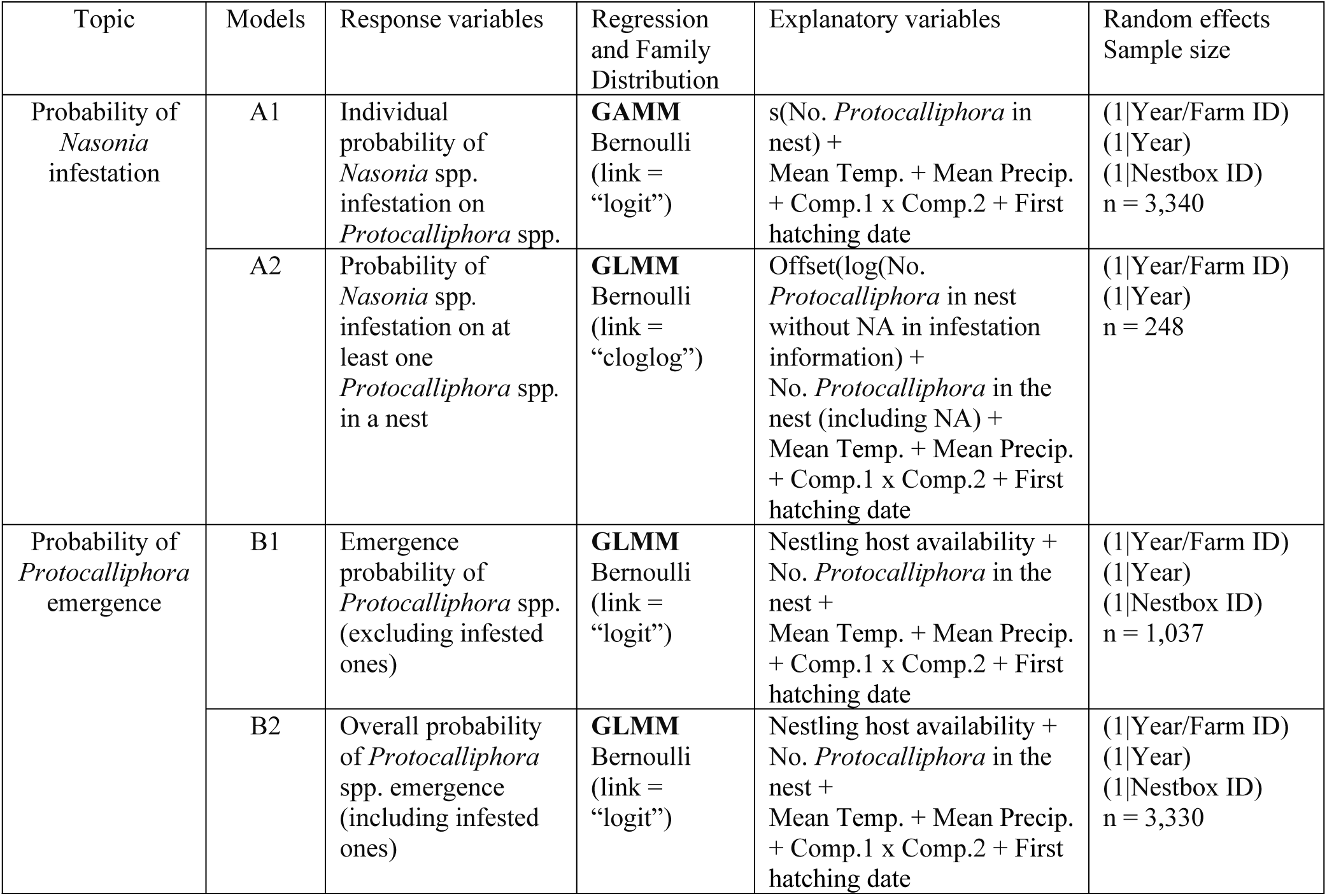
Description of the different models built according to Directed Acyclic Graph (DAG) with respective family distribution and sample size. Comp.1 & Comp.2 : Components of the robust Principal Component Analysis. Temp. & Precip. : Temperature and precipitation (averaged during either nestling or *Protocalliphora’s* development period.

Weather conditions during the larval/pupation phases corresponded to hourly averages over the respective period during which swallow nestlings and blowfly larvae and pupae developed in each swallow brood (i.e., up to fledging or death and to estimated emergence, respectively).

Agricultural context surrounding each swallow brood was defined based on the scores of the first two axes of a robust, compositional PCA fitted on relative land cover values measured annually within 500 m of each nest box of the study system (i.e., Comp.1 and Comp.2 in Fig. 5A).

We fitted four Bayesian models, either General Additive Mixed Models (GAMMs) or General Liner Mixed Models (GLMMs) as described in Table 1, using the *brms* package (v2.21; Bürkner 2017) in R (v.4.3.1, R core teams 2023) with non- or weakly informative priors. We derived posterior distributions with four independent Markov Chain Monte Carlo (MCMC). We extracted posterior draws with *tidybayes* package (v3.0.6; Kay 2023), with respective credibility intervals at 50-80-95%. We then extracted Conditional and Marginal R² with the function r2_bayes from the *performance* package (v0.10.8; Lüdeck *et al*. 2021).

The first two models (A1 and A2, Table 1), explain the variation in the probability of *Protocalliphora* infestation by *Nasonia*. Model A1 considers the likelihood of recording at least one parasitoidism event within a nest containing *Protocalliphora*. Model A2 examines the proportion of *Protocalliphora* infested by *Nasonia* within a given nest. The other two models (B1 and B2, Table 1) concern *Protocalliphora* emergence probability. Model B1 excludes *Nasonia* parasitoidism, focusing only on the effects of other probable causes of non-emergence, such as direct agricultural activity impacts. Model B2 includes all *Protocalliphora*, regardless of infestation, without controlling for “infestation effects on emergence success” (Fig. 2). This approach helps to understand the total effect of agricultural landscapes on *Protocalliphora* emergence at the population scale, which can potentially be driven indirectly by landscape impacts on *Nasonia* parasitoidism. Given the hierarchical structure of our dataset, we accounted for non-independence in the data using random effects. Nestbox ID was included as a random intercept to control for micro-environmental variation at the nest scale (e.g., vegetation, microclimate) as well as potential parental effects, since the same adult *Protocalliphora* or *Nasonia* individuals may have laid multiple eggs within a nestbox. We also included Farm ID nested within Year as a random effect. This structure captures farm-specific effects not fully explained by our environmental variables (e.g., pesticide use, management style), which may vary not only between farms but also across years within the same farm. Finally, Year was added as a separate random effect to account for broader interannual variability in ecological conditions (e.g., weather, phenology) that were not explicitly measured in the dataset.

## Results

### Density-dependence and weather influence *Nasonia* infestations

The proportion of *Protocalliphora* infested by *Nasonia* across all subsampled nests was 68.65% (2293 of 3340), leaving 31.35% uninfested (1047). These *Protocalliphora* originated from 248 distinct nests, 211 (85.80%) of which counted at least one *Nasonia* parasitism event. The number of *Protocalliphora* individuals within a nest influenced the probability of infestation by *Nasonia* both at the individual level (model A1; R²c = 23.7%; R²m = 11.7%; Fig. 3A) and at the scale of the nests (model A2; R²c = 54.4%, R²m = 34.2%; Fig. 3A). As the number of *Protocalliphora* in a nest increased, so did the likelihood of an infestation at the nest level. Model predictions suggested nests containing more than 25 *Protocalliphora* have more than a 95% probability of *Nasonia* parasitoidism occurrence. At the individual level, the infestation probability rose from 0.55 to 0.75 as the number of *Protocalliphora* in a nest increased from 1 to 20. However, as the number of *Protocalliphora* continued to rise, a dilution effect was observed: the infestation probability per individual decreased, dropping to 15% in nests with ∼ 100 hosts. Theoretically, the infestation probability predicted by both models at x=1 (a single *Protocalliphora* within a nest) should be the same, as the likelihood of a nest containing at least one infestation is equivalent to the probability of that single *Protocalliphora* being infested. The slight discrepancy in intercepts at x=1 in Fig. 3A, arises from the model structure: one is a GAMM with a logit link function analyzing 3,340 observed individuals across 248 nests) while the other is a GLMM with a cloglog link applied to the nest-level scale, focusing on 248 nest observations.

**Figure 3.**
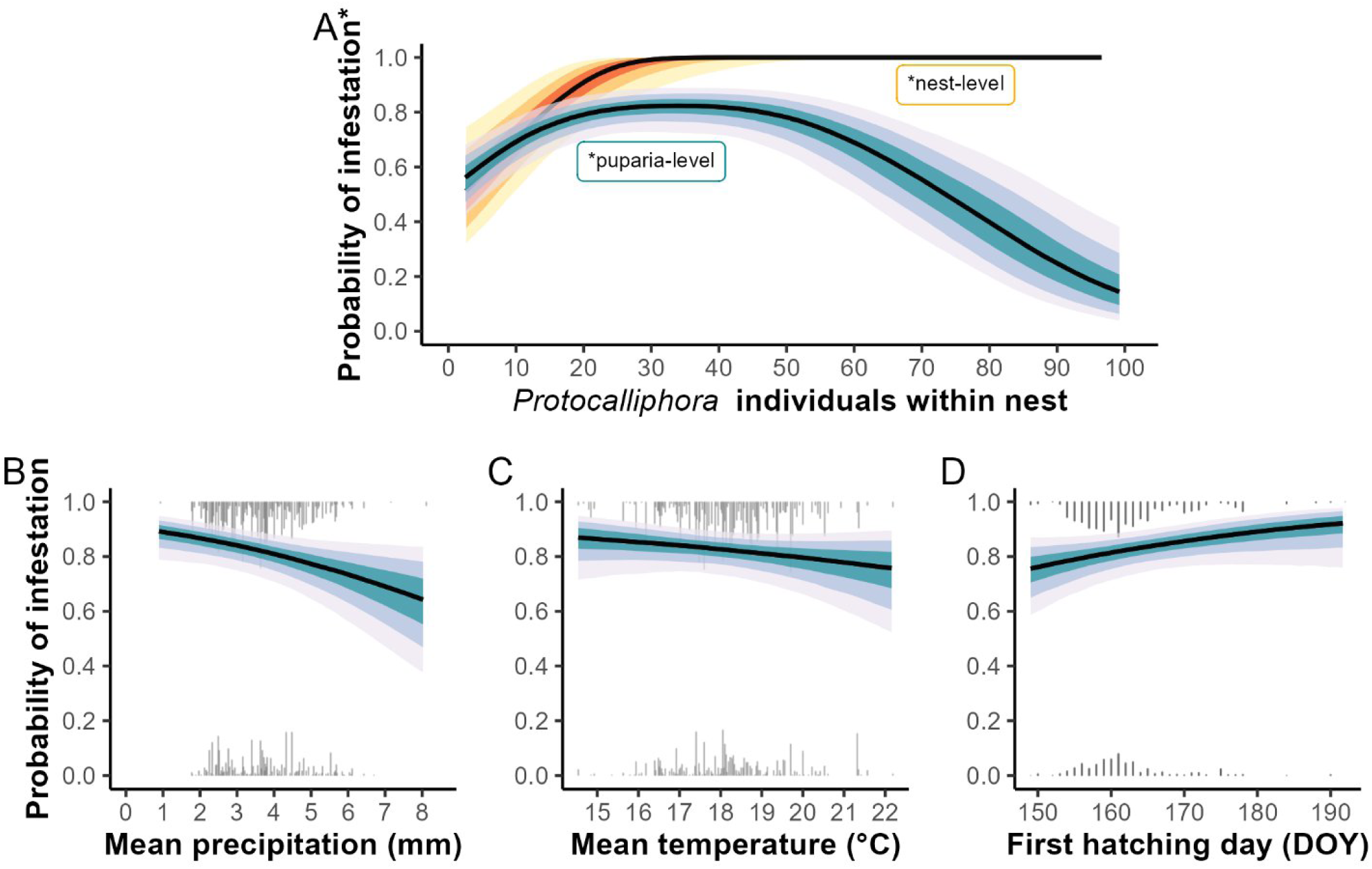
Infestation probability by *Nasonia* spp. at both the scale of nests infested by *Protocalliphora* (orange, n = 248) and the individual *Protocalliphora* scale (blue, n = 3340), with 50%,80% and 95% credibility intervals, according to: A) Number of *Protocalliphora* hosts within the nest, B) precipitation (mm) during puparia development, measured from the first Tree swallow nestling hatching to 7 days following the last fledge or death, C) Temperature (°C) during puparia development, measured from the first Tree swallow nestling hatching to 7 days following the last fledge or death, D) timing (Day of year) of the first Tree Swallow hatching in a clutch containing *Protocalliphora* spp.

Both temperature and precipitation during the *Protocalliphora* development and pupation periods (from the first avian nestling hatch event to seven days after the last nestling fledges or dies) emerged as direct causal factors for individual infestation probability. Both higher rainfall and higher temperature decreased the likelihood of *Nasonia* infestation by about 20%, from the lowest to the highest observed values of these weather variables (Fig. 3B, C). *Protocalliphora* individuals infesting later Tree Swallow clutches were more likely to be infested by *Nasonia* parasitoid wasps (90% in mid-July) than those infesting earlier-laid clutches (75% at the end of May ; Fig. 3D).

### Nestling number and weather influence *Protocalliphora* emergence

Approximately 68.7% of *Protocalliphora* puparia were infested by *Nasonia* parasitoid wasps. Among the 1037 uninfested individuals, 624 (60.17%) emerged, while 413 (39.83%) failed to emerge due to reasons other than *Nasonia* infestations. Model predictions indicated that *Protocalliphora* probability of emergence increased steeply with a greater number of nestling hosts for a longer period (Fig. 4A), rising from 0.22 to 0.74 from the lowest to the highest observed values of nestling availability. The probability of emergence success in uninfested individuals also increased with higher average temperatures during the *Protocalliphora* development period (Fig. 4B), ranging from 0.29 to 0.83 across the observed range of average temperature values. Conversely, both the abundance of *Protocalliphora* within the nest (Fig. 4C) and later phenology of the bird host (Fig. 4D) were negatively associated with the probability of emergence, when considering only uninfested individuals. For instance, a solitary *Protocalliphora* individual in a Tree Swallow nest had an estimated emergence probability of around 62%, whereas this probability dropped to 49% in the presence of 99 competitors. Regarding phenology, although few first clutches were initiated after day 175 of the year (Doy), emergence probability declined markedly with seasonal progression, falling from 0.78 at Doy 150 to just 0.32 by Doy 175.

**Figure 4.**
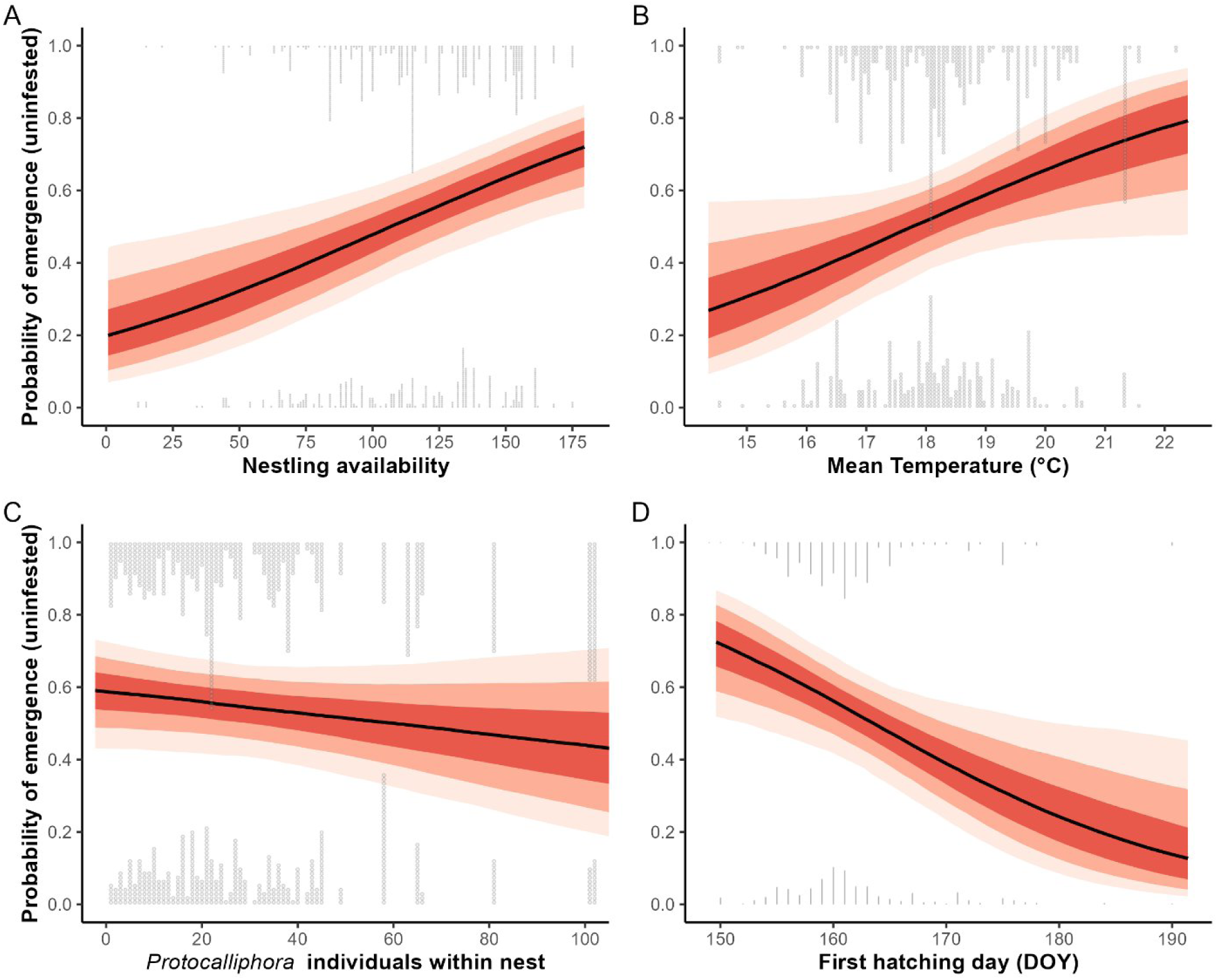
Predicted mean emergence probability of *Protocalliphora* not infested by *Nasonia* spp., with 50%,80% and 95% credibility intervals, according to A) the number of nestling-days within the nest, B) Temperature (°C) during puparia development, measured from the first Tree swallow nestling hatching to 7 days following the last fledge or death, C) the abundance of *Protocalliphora* within the nest D) timing (Day of year) of the first Tree Swallow hatching in a clutch containing *Protocalliphora* spp.

### Agricultural contexts influence *Protocalliphora* emergence and *Nasonia* infestation

The Robust Principal Components Analysis (rPCA) and the yearly landscape characterization for each nestbox show that component 1 (Comp1) correlated positively with corn and soybean fields and negatively with forage and tree cover, explaining approximately 78.98% of habitat variance (Fig. 5A). Component 2 (Comp2) correlated positively with tree cover and negatively with forage, explaining about 16.68% of the variance. Grey dots (Fig. 5A) represent agricultural contexts within a 500m radius of all nest boxes, while darker dots represent those in which we analyzed *Protocalliphora* individuals and obtained data on either infestation or emergence probability. Connecting outermost dots delineates the boundaries of a polygon that encompasses the full range of observable agricultural landscapes within our study system (Fig. 5A).

**Figure 5.**
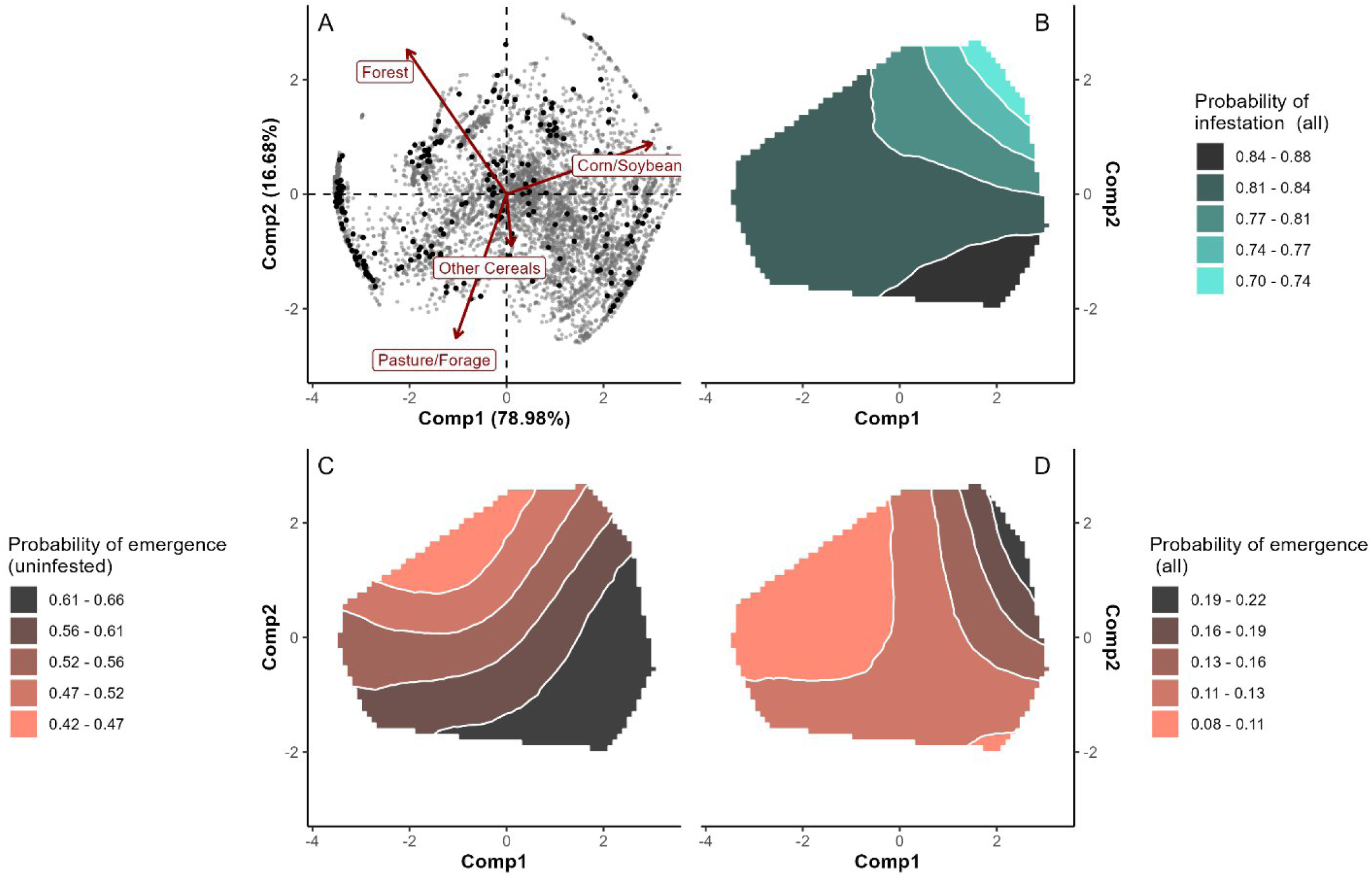
A) Robust Principal Components Analysis (rPCA) of the landscape within a 500m radius surrounding the 400 nestboxes in the study system between 2006-2019 (n = 5,600) B) Infestation probability of all *Protocalliphora* by *Nasonia* spp. (n = 3,340) C) Emergence rate of uninfested *Protocalliphora* (n = 1,037) D) Emergence of all *Protocalliphora* (including infested ones; n = 3330). Darker dots in Fig. 5A represent the agricultural contexts surrounding the nests containing *Protocalliphora* puparia subsampled for this study (n = 248). Other figures represent heatmaps of the effects of agricultural contexts, approximated via the interaction of Comp1 and Comp2 The left side of the polygon encompasses landscapes dominated by agroextensive practices, such as pastures, forages with more tree cover in the top-left corner while the right side represents agrointensive practices, primarily featuring corn and soybean fields with minimal tree cover in the top-right corner, and mixed cereal crops with little to no tree cover in the bottom-right corner.

Agricultural contexts emerged as factors influencing substantially both the probability of *Protocalliphora* infestation by *Nasonia* and the probability of *Protocalliphora* emergence. The probability of *Nasonia* infestation in *Protocalliphora* individuals (predicted by model A1) decreased from around 0.88 in agroextensive contexts to 0.70 in agrointensive monocultures of corn and soybean (Fig. 5B). The emergence probability of uninfested *Protocalliphora* (Model B1; Fig. 5C) rose from 0.42 in agroextensive contexts with high tree cover to 0.66 in agrointensive areas with a mix of corn, soybean, and other cereals, as well as pastures and forages with low tree cover (low Comp. 2 values).

As agricultural context affects *Protocalliphora* emergence both indirectly (through infestation probability; Fig. 5B) and directly (Fig. 5C), Model B2 examined the total effects of agricultural practices on the emergence probability of all *Protocalliphora*, including failures induced by *Nasonia* ranging from 0.08 to 0.22 (Fig. 5D). Overall, emergence probability for all *Protocalliphora* was inversely related to trends of *Nasonia* infestation probability. Lower global probabilities of emergence were observed in agroextensive contexts with high tree cover, where *Nasonia* infestation probability was high (Fig. 5B). In these areas, emergence probability was also low for uninfested *Protocalliphora* (Fig. 5C). The probability of emergence for all *Protocalliphora* gradually increased to a maximum in agrointensive contexts with high tree cover, where *Nasonia* infestation probability was reduced (Fig. 5B).

## Discussion

The understanding of density-dependent mechanisms is crucial for grasping the trophic relationships that govern host-parasite networks (Herrando-Pérez *et al*. 2012, Lessells 1985). In this study, we investigated how density-dependent mechanisms and agroenvironmental factors influence infestation and survival rates within a tri-trophic system comprising Tree Swallow nestlings, their nest-dwelling ectoparasites (*Protocalliphora*) and the parasitoid jewel wasps (*Nasonia*). Anthropogenic disturbances do not simply reduce biodiversity but can fundamentally reorganize interaction networks through density-dependent and indirect pathways.

### Density-dependence and nest characteristics affect infestation and emergence probability

A study has shown that the number of Tree Swallow nestlings-days increased both the probability of occurrence and the abundance of *Protocalliphora* ectoparasites (Coroller-Chouraki *et al*. 2026b). Corroborating these findings, the present study found that higher nestling availability also positively affected *Protocalliphora* emergence success, provided that the pupae were not infested by *Nasonia* parasitoids. This result supports the suggestion that more food availability for an extended period prevents *Protocalliphora* from being underfed and from pupating prematurely as “runts” (Sabrosky *et al*. 1989), potentially leading to death during pupation. Controlling for avian host density, we found that higher abundances of *Protocalliphora* within nests slightly reduced the emergence success of individuals that were not parasitized by *Nasonia,* suggesting the possibility of intra- and interspecific competition among *Protocalliphora* spp., potentially due to overcrowding on limited accessible areas of the nestlings bodies, such as the legs, which may restrict feeding opportunities for weaker larvae.

*Protocalliphora* emergence success can be significantly influenced by agroenvironmental factors, with the primary factor being *Nasonia* parasitoid infestation. These wasps effectively control insect host populations (Werren and Loehlin 2009) and in our sample, 68.65% of the *Protocalliphora* puparia either contained *Nasonia* pupae or were empty with *Nasonia* exit holes, indicating that the *Protocalliphora* were devoured by *Nasonia*. The likelihood of *Nasonia* parasitizing at least one *Protocalliphora* in each Tree Swallow nest increased sharply with the number of *Protocalliphora* present. A study by Garrido-Bautista *et al*. (2020) in central and southern Spain also found a positive association between the number of *Protocalliphora* and prevalence of parasitoidism in nests of pied flycatcher (*Ficedula hypoleuca*) and blue tits (*Cyanistes caeruleus*). Our model predicted that nests with more than 10 *Protocalliphora* systematically experience at least one *Nasonia* parasitoidism event. Besides the mechanistic higher probability of an event occurring with an increased number of trials, this phenomenon could more likely be explained by the fact that a higher number of *Protocalliphora* hosts can potentially lead to increased olfactory stimuli that *Nasonia* wasps use to locate and parasitize them (Frederickx *et al*. 2013). Additionally, a larger number of *Protocalliphora* hosts, potentially originating from different oviposition events, might extend the time window of their availability for *Nasonia* infestation. A greater number of Tree Swallow nestlings developing over an extended period in the nest resulted in higher *Protocalliphora* ectoparasite loads (Coroller-Chouraki *et al*. 2026b). As a result, elevated *Protocalliphora* densities in a given site and year reflect increased host availability for *Nasonia* parasitoid wasps, potentially allowing *Nasonia* populations to surge the following year. However, no study to date has explicitly explored whether these two insect taxa follow cyclical, predator-prey-like dynamics akin to Lotka-Volterra models.

While nests with greater abundance of *Protocalliphora* hosts inevitably experience *Nasonia* infestations, the proportion of infested *Protocalliphora* hosts in each nest initially rises, then plateaus, and finally decreases, indicating a strong dilution effect. Although multiple *Nasonia* individuals can lay eggs in the same *Protocalliphora* host (Godfray 2010), a high number of potential hosts might trigger host-selection and host-rejection mechanisms by *Nasonia*. *In-vitro* studies have shown that the number of eggs laid, and the offspring sex ratio, depend on the host’s potential quality, on whether another *Nasonia* had previously oviposited within the host, and on how long ago that event occurred (Koppik *et al*. 2019, 2014, Shuker *et al*. 2005). However, *Nasonia’s* behaviour during oviposition is not well understood. Similar density-dependent parasitoid responses have also been observed in cavity-nesting Hymenoptera systems where Wild et al. (2026) found that parasitism by *Melittobia* spp. increased with host availability across agricultural landscapes, while landscape context differently affected occupancy and reproductive performance among trophic groups. These results further support the idea that agroenvironmental conditions can restructure multitrophic interactions through context-dependent pathways.

In natural conditions, individual *Nasonia* may leave an avian nest without ovipositing in any *Protocalliphora* puparia and search for another host patch after rejecting hosts that were already infested by conspecific individuals (Gunton and Pöyry 2016, Koppik *et al*. 2019, Shuker *et al*. 2005). Combined with differences in dispersal ability and reproductive rate between parasitoids and their hosts, such selective foraging behavior can lead to a functional saturation threshold and to an under-aggregation of parasitoids within dense host patches (Comins and Hassell 1979, Gunton and Pöyry 2016, Hassell 2000). This density-dependent nonlinearity can generate apparent dilution effects, as the proportion of infested hosts declines despite their higher abundance (Gunton and Pöyry 2016, Lessells 1985, Otway *et al*. 2005). Similar patterns have been described in many different systems, from nematodes to dipteran parasitoids to aquatic trematodes, illustrating how behavioral constraints and resource heterogeneity shape the strength of trophic interactions and trophic networks stability (Gunton and Pöyry 2016, Morand *et al*. 1996).

In our system, the probability of *Protocalliphora* being parasitized by *Nasonia* wasps increased slightly with later hatching dates of Tree Swallow clutches. This increase reflects the multivoltine nature of *Nasonia* wasps (Paolucci et al. 2013). As the breeding season progresses, the emergence of successive wasp generations potentially leads to a higher density of gravid females available to oviposit in late-season nests. Furthermore, *Protocalliphora* in later Tree Swallow clutches were also much less likely to emerge, even if uninfested. Merino et al. (2024) reported comparable results, potentially due to the advanced physiological age of ovipositing females later in the breeding season, which could alter offspring quality. High densities of *Protocalliphora* within a given nest were associated with lower probability of emergence for individuals not infested by *Nasonia*. Future studies could examine whether competition does affect different *Protocalliphora* species equally within the same nest.

### Effects of weather on infestation and emergence probability

Both higher temperature and precipitation during *Protocalliphora* development slightly decreased their probability of being infested by *Nasonia*. This result differs from Garrido-Bautista *et al*. (2020)’s finding, where moister habitat increased *Nasonia* infestation prevalence. Our results suggest that *Nasonia* movements and host searching during oviposition could be hindered by heavy rain or high humidity; as for the effects of higher temperatures, several physiological and behavioral hypotheses can explain the lower infestation probability. For instance, *Nasonia* may adopt avoidance strategies or enter dormancy during heatwaves, or experience reduced fecundity, similarly to other insects (Ratte 1985, Zhang *et al*. 2013), including some parasitoid wasps (Bressac *et al*. 2023). The literature about the effect of temperature on *Nasonia* mostly consists of *in-vitro* experiments and most studies focus on the developmental period (Hidalgo *et al*. 2019, Zhang *et al*. 2013, 2019), obscuring the effect of temperature in natural conditions. Higher temperatures could have indirect effects on *Nasonia* infestations by reducing the time required for its pupation, decreasing *Nasonia*’s ability to locate and infest the pupae. Indeed, Bennett and Whitworth (1991) showed that, like other *Protocalliphora* species, *P. sialia* pupates faster in warmer conditions, averaging 58 hours at 35°C compared to 355 hours at 13°C. These authors also reported that, up to a degree, warmer conditions led to substantially higher emergence probabilities in uninfested individuals (Dawson *et al*. 2005). The fact that warmer conditions exert opposing pressures on this multi-trophic system, by reducing *Nasonia* parasitism while enhancing the abundance and emergence probability of its host, underscores why weather variables must be integrated into complex interaction network analyses (Thierry et al. 2019).

### Effects of agricultural intensification on infestation and emergence probability

When not infested, *Protocalliphora* tended to emerge more frequently in areas with fewer trees, both in agrointensive and agroextensive contexts. This pattern contrasts with expectations based on habitat quality (Coroller-Chouraki *et al*. 2026b), suggesting a potential survival bias (Zens and Peart 2003). *Protocalliphora* ectoparasites are counted after nests collection only if they have successfully pupated, as larvae that died before pupating are not detected. Early death of low-quality larvae in harsher habitats, such as agrointensive landscapes (Raven and Wagner 2021), could lead to the survival of only the fittest individuals within nests located in such agricultural contexts. A more pronounced counter-selection process in agrointensive compared to agroextensive landscapes could therefore explain why the sampling of *Protocalliphora* puparia in our system reveals a reduced abundance (Coroller-Chouraki *et al*. 2026b) but a higher emergence probability of uninfested individuals within agrointensive landscapes, as only the fittest larvae would survive to pupation.

The agricultural landscape significantly influences the probability of *Protocalliphora* being parasitized by *Nasonia* wasps, as in areas dominated by corn and soybean monocultures with low tree cover (characteristically agrointensive landscapes) the likelihood of *Nasonia* infestation events was the lowest. This probability increases as the habitat shifts away from such monocultures. One plausible explanation for the lower *Nasonia* infestation in agrointensive landscapes would be the result of a trophic cascade, with lower avian host availability, lower *Protocalliphora* ectoparasite abundance, and lower parasitism infestation rate, due to spill-over effects in such contexts (Daoust *et al*. 2012a, Daszak *et al*. 2000, Wells *et al*. 2015). Alternatively, agrointensive contexts, where pesticides are more prevalent and abundant (Giroux 2022, Poisson *et al*. 2021) could reduce *Nasonia’s* ability to find sexual partners and hosts, as suggested by *in-vitro* studies (Schöfer *et al*. 2023, Sulg *et al*. 2023), leading to lower infestation rates. *Nasonia* survival, body condition, fecundity, and offspring quality could also be altered by pesticide exposure, including potential cocktails effects in agrointensive contexts (Pisa *et al*. 2015, Sanchez-Bayo and Goka 2014). A less likely alternative explanation for these results is that *Nasonia* wasps exhibit plasticity in their host-selection behavior in agrointensive landscapes, potentially neglecting their primary *Protocalliphora* hosts which are less abundant in these habitats.

The significant role of *Nasonia* infestation in controlling *Protocalliphora* populations and its notably lower infestation rate in agrointensive habitats influences overall *Protocalliphora* emergence. In agrointensive landscapes where *Nasonia* infestation probability is minimal, there is a higher likelihood of *Protocalliphora* emergence, despite *Protocalliphora* being less abundant in such contexts (Coroller-Chouraki *et al*. 2026b). We used agroextensive landscapes as a control habitat to assess the effects of agricultural intensification on ectoparasite loads. However, this argument must be contextualized as some pesticides used in pastures (considered as agroextensive landscapes) may also negatively impact the entomofauna (Buijs *et al*. 2022).

“Our study demonstrates that agrointensive landscapes suppress not only avian hosts, but also their *Protocalliphora* ectoparasites and associated *Nasonia* parasitoids. While the resulting release from top-down control seemingly boosts *Protocalliphora* emergence, this trend could be driven by a survival bias, where unfit individuals are counter-selected early on in high-stress habitats. Sampling larvae before pupation will be essential for future protocols to untangle these hidden dynamics. This methodological nuance underscores how vulnerable intricate, unmanaged trophic networks are to being disrupted (or wiped out) by agricultural practices before we can fully understand them. While literature frequently details impacts on crop-related species, our findings bridge a critical gap regarding how landscape modification alters non-target biological control (Gagic et al. 2012). Ultimately, this work provides strong evidence that intensive agriculture, through both landscape homogenization (Kovács-Hostyánszki et al. 2017) and pesticide pressure (Uhl and Brühl 2019), can adversely affect non-targeted and non-crop-related insect communities.

## Acknowledgements

The authors extend their deepest gratitude to the farm owners whose cooperation was vital to our long-term monitoring program. We also thank the Université de Sherbrooke for its institutional support and the funding agencies that facilitated this research, specifically the Natural Sciences and Engineering Research Council of Canada (NSERC), the Fonds de recherche du Québec – Nature et technologies (FRQ-NT), and the Canada Research Chairs program. We are indebted to the many students, field assistants, and technicians involved in the study, with a special mention to L. Glaude and Y. Sageau for their dedicated work in nest collection and processing over the years. We thank T. Whitworth for his invaluable taxonomic expertise regarding challenging *Protocalliphora* specimens, and D. Garneau for his generous support with imagery. This study was conducted under the approval of the Université de Sherbrooke’s animal care committee, and we gratefully recognize everyone whose efforts, both in the field and behind the scenes, made this work possible.

