## Supplementary material for "Density-Dependence and agricultural intensity affect host-parasite-parasitoid interactions in a bird-insect trophic system": Fig. S1

-
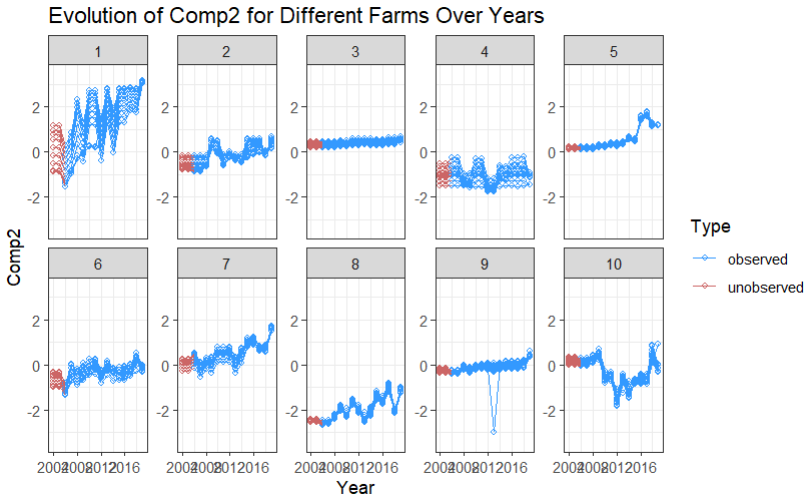

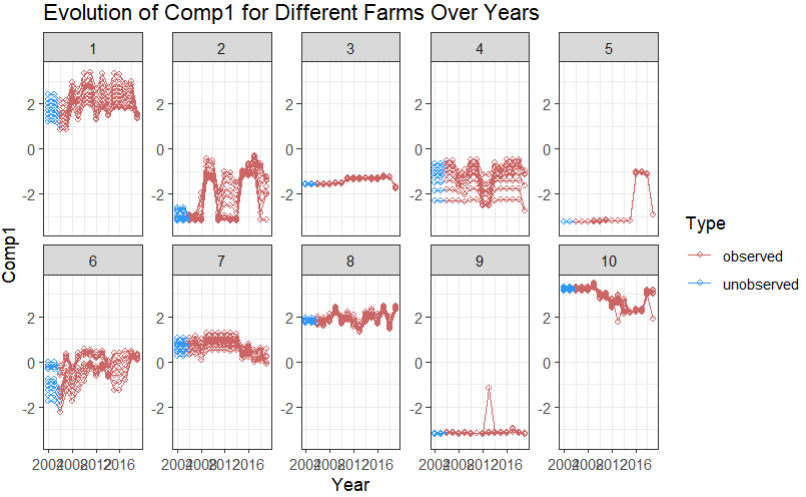
**Extrapolation of 2004 and 2005’s Comp.1 and Comp.2 values**

**Figure S1.** Interannual stability of landscape composition around Tree Swallow nest boxes between 2004 and 2019. Extrapolated Comp.1 or Comp.2 for 2004 and 2005 missing values are displayed in contrasting color (respectively blue vs red, and red vs blue).
